## Supporting Information for "High Resolution Biomolecular Condensate Phase Diagrams with a Combinatorial Microdroplet Platform"

### Supporting Information: High Resolution and Multidimensional Protein Condensate Phase Diagrams with a Combinatorial Microdroplet Platform

William E. Arter,[1,†] Runzhang Qi,[1,†] Nadia A. Erkamp,[1,†] Georg Krainer,[1,†] Kieran Didi,[1] Timothy J. Welsh,[1] Julia Acker,[2] Jonathan Nixon-Abell,[3] Seema Qamar,[3] Jordina Guillén-Boixet,[4] Titus M. Franzmann,[4] David Kuster,[5] Anthony A. Hyman,[5] Alexander Borodavka,[2] Peter St George-Hyslop,[3,6,7] Simon Alberti[4] and Tuomas P.J. Knowles[1,8,*]

*[1] Yusuf Hamied Department of Chemistry, Centre for Misfolding Diseases, University of Cambridge, Lensfield Road, Cambridge, CB2 1EW, UK*

*2 Department of Biochemistry, University of Cambridge, Cambridge, CB2 1QW, UK*

*3Cambridge Institute for Medical Research, Department of Clinical Neurosciences, University of Cambridge, Cambridge CB2 0XY, UK*

*[4] Biotechnology Center (BIOTEC), Center for Molecular and Cellular Bioengineering (CMCB), Technische Universität Dresden, Tatzberg 47/49, 01307 Dresden, Germany*

*[5]Max Planck Institute for Molecular Cell Biology and Genetics, Pfotenhauerstrasse 108, 01307 Dresden, Germany*

*6 Department of Medicine (Division of Neurology), University of Toronto and University Health Network, Toronto, Ontario M5S 3H2, Canada*

*[7]Department of Neurology, Columbia University, 630 West 168th St , New York, NY 10032, USA.*

*[8] Cavendish Laboratory, Department of Physics, University of Cambridge, J J Thomson Ave, Cambridge, CB3 0HE, UK*

*† These authors contributed equally.*

* To whom correspondence should be addressed:


#### Supporting Results

##### Fluorescence recovery after photobleaching of FUS$^{G156E}$-EGFP condensates

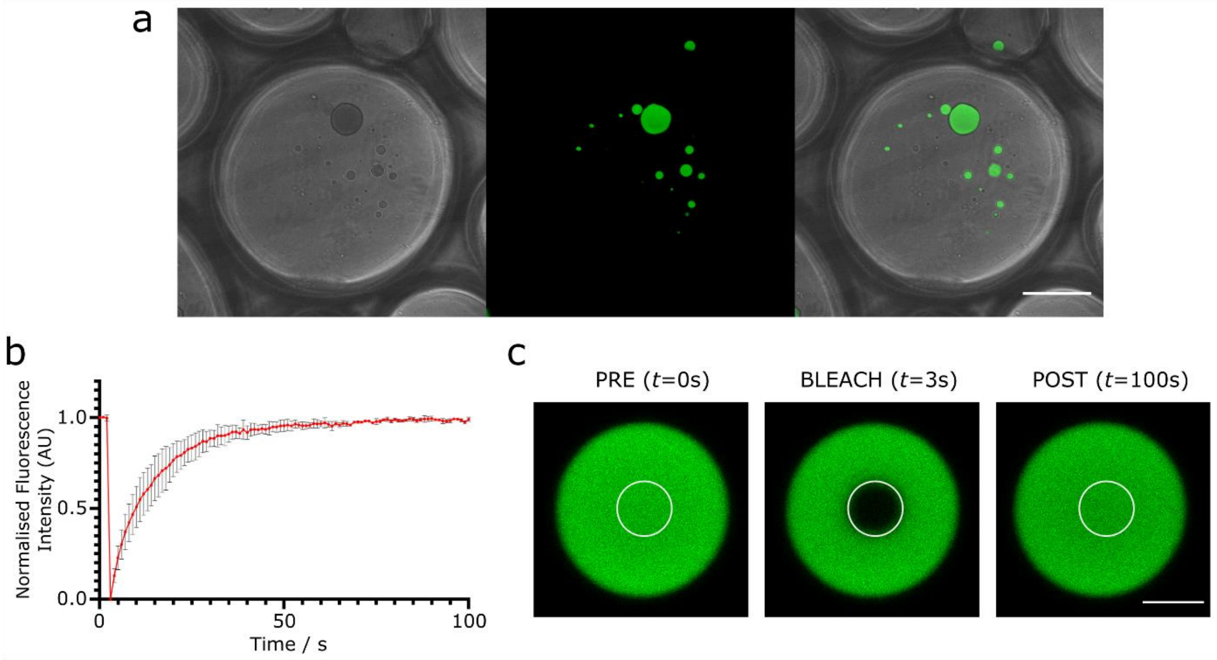


**Figure S1. Fluorescence recovery after photobleaching of FUS$^{G156E}$-EGFP condensates. (a)** Brightfield (left), fluorescence (centre), and merged (right) fluorescence microscopy images of water-in-oil droplets containing FUS$^{G156E}$-EGFP condensates. Scale bar 50 µm. **(b)** Fluorescence recovery data for FUS$^{G156E}$-EGFP condensates encapsulated in water-in-oil droplets. Error bars correspond to standard deviation of three FRAP acquisitions of different condensates. Scale bar 5 µm.. **(c)** Exemplary FRAP data before (PRE, $t$ = 0s), shortly after photo bleaching pulse (BLEACH, $t$ = 3s), and after fluorescence recovery (POST, $t$ = 100s).

The liquid nature of homotypic FUS condensates (4 µM FUS$^{G156E}$-EGFP, 4% *w/v* PEG 6k) encapsulated within oil-in-water microdroplets was confirmed by fluorescence recovery after photobleaching (FRAP) experiment (Figure S1). Fluorescence recovery occurred within 50 ms (mobile fraction 98.3 ± 0.63 %), indicating that condensates retained their liquid nature when encapsulated within fluorinated oil in the presence of 1.5% fluorosurfactant.

#### Comparison between PhaseScan and manual measurements

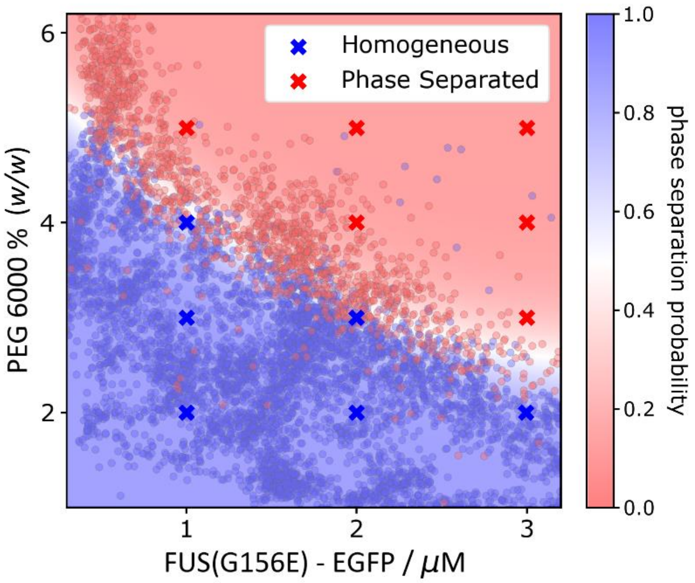


**Figure S2. Comparison between PhaseScan and manual measurements.** The crowding-induced homotypic phase separation of EGFP-tagged $FUS^{G156E}$ in the presence of PEG 6000 was analysed by independent PhaseScan (dots and colour map) and manual pipetting experiments (crosses). All points were observed to match the phase diagram as determined by PhaseScan.

To verify that the phase diagram generated by the PhaseScan platform is in accordance with the phase diagram obtained from bulk experiments, we performed manual pipetting experiments with $FUS^{G156E}$ and compared the phase behaviour with the data obtained by PhaseScan (Figure S2). We performed manual bulk measurements at twelve different points in the phase diagram, stock solutions of buffer, 6k PEG (20%) and FUS (18.23 μM as verified by Nanodrop measurements at 488 nm) were prepared and mixed to give 10 μL of each of the sample compositions as shown in Figure S2. The sample was pipetted onto a microscope slide equipped with imaging chambers, which was sealed with a coverslip, before imaging EGFP fluorescence with a 10× objective. The images were manually classified as containing phase-separated or homogeneous protein and compared to the corresponding region in the phase diagram produced by the PhaseScan platform. All manually determined datapoints match well with the boundary as determined by PhaseScan platform.

#### Effect of barcoding dyes on phase behaviour

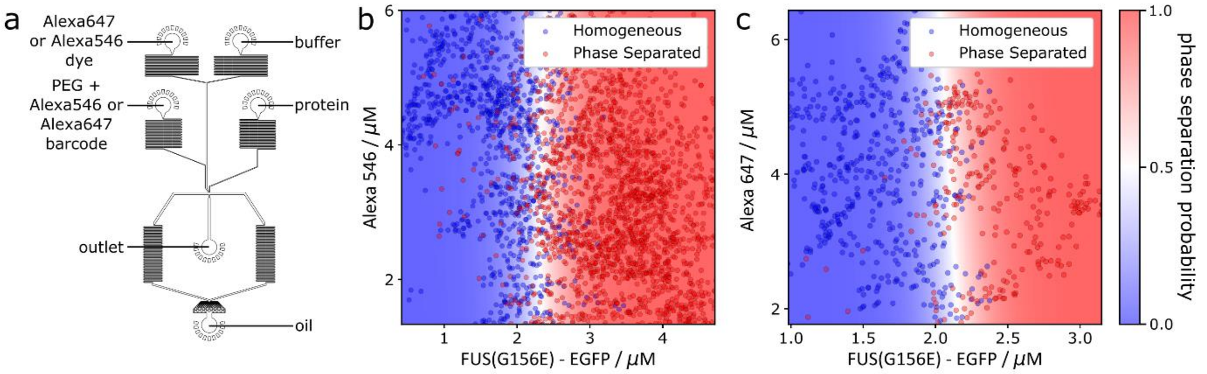


**Figure S3. Effect of barcoding dyes on phase behaviour. (a)** PhaseScan chip used to assess dye behaviour. **(b)** Phase diagram of EGFP-tagged FUS$^{G156E}$ vs. increasing concentrations of the barcoding dye Alexa 546. **(c)** Phase diagram of EGFP-tagged FUS$^{G156E}$ vs. increasing concentrations of the barcoding dye Alexa 647.

To assess whether the fluorescent dyes used to barcode for different PhaseScan solution components effect phase behaviour, control experiments were performed in which EGFP-FUS$^{G156E}$ protein and barcoding dye concentrations were varied in the presence of a single concentration (2.4% (*w*/*w*)) of PEG 6000. This was achieved through the use of a 4-variable droplet generator as shown in Figure S3(a). We introduced Alexa647 or Alexa546 dye, buffer and protein solutions into the chip with varying flow rates according to an automated flow programme (total flow rate 60 μL/h) with the Alexa546 or Alexa647-barcoded PEG solution injected into the chip at a rate of 7 μL/h. Droplets were collected and analysed as described in the Methods. Data points possessing anomalous PEG barcode concentrations (due to variance in the PEG flow rate) were removed, and the resultant phase diagrams for EGFP-FUS$^{G156E}$ concentration vs. Alexa546 or Alexa647 concentrations were produced (Figure S3(b, c)), respectively). We observed no significant variation in the position of the phase boundary with respect to Alexa546 or Alexa647 dye concentration, indicating that the phase behaviour is independent of barcode dye concentration.

#### Time dependence, reproducibility, and minimal effect of outliers

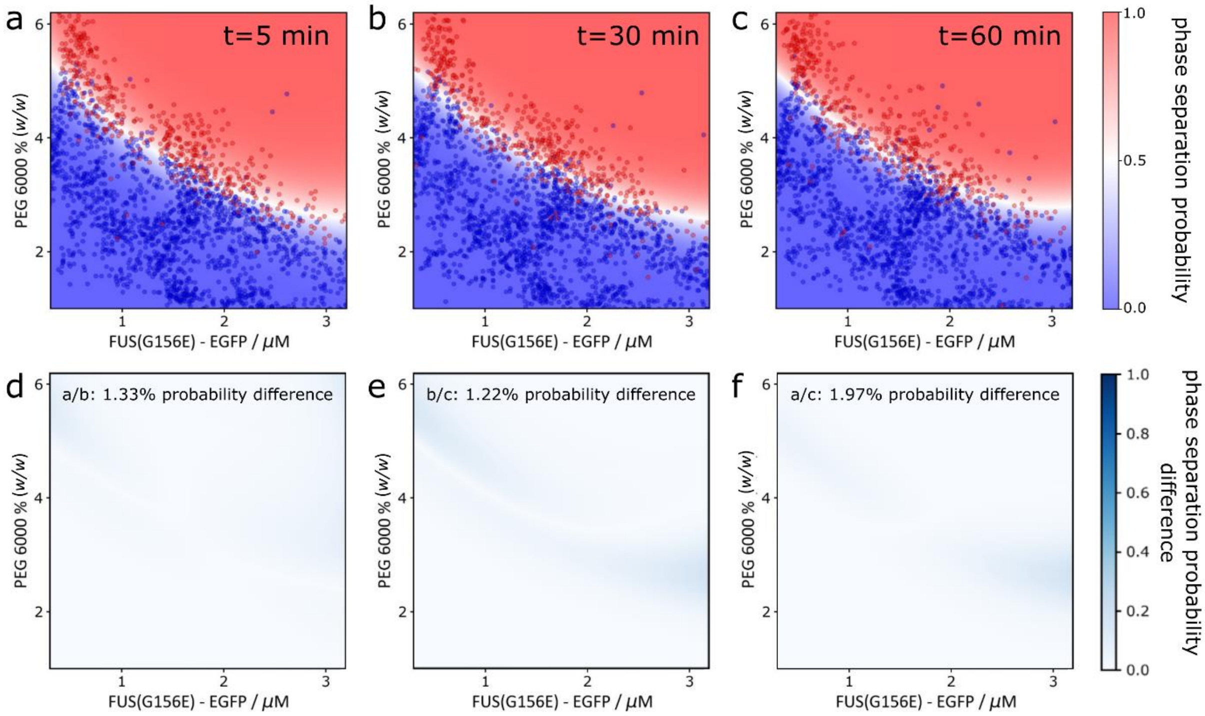


**Figure S4. Reproducibility of PhaseScan method, minimal effect of outliers, and time-dependence of phasescan measurement. (a–c)** Phase diagrams corresponding to separate phase scan experiments repeated on the same protein solutions, data collection was undertaken 5 min after droplet generation for (a), 30 min for (b) and 1 h for (c). **(d–f)** Plots comparing reported phase separation probability in experiments panels a–c. Only negligible differences are observed in the integrated phase separation probability between the repeats (average difference 1.51%), demonstrating reproducibility of droplet production, trapping, data acquisition and analysis. Notably, although a small number of incorrectly classified or barcoded outliers exist in all three datasets, these datapoints have negligible effect on the reported position of the phase boundary since they exist as only a small (< 2%) proportion of the overall dataset. The negligible differences between the phase diagrams shows that there is no time-dependence in the observed phase behaviour, confirming that the phase diagrams are collected under equilibrium conditions.

#### Effect of droplet size on phase behaviour

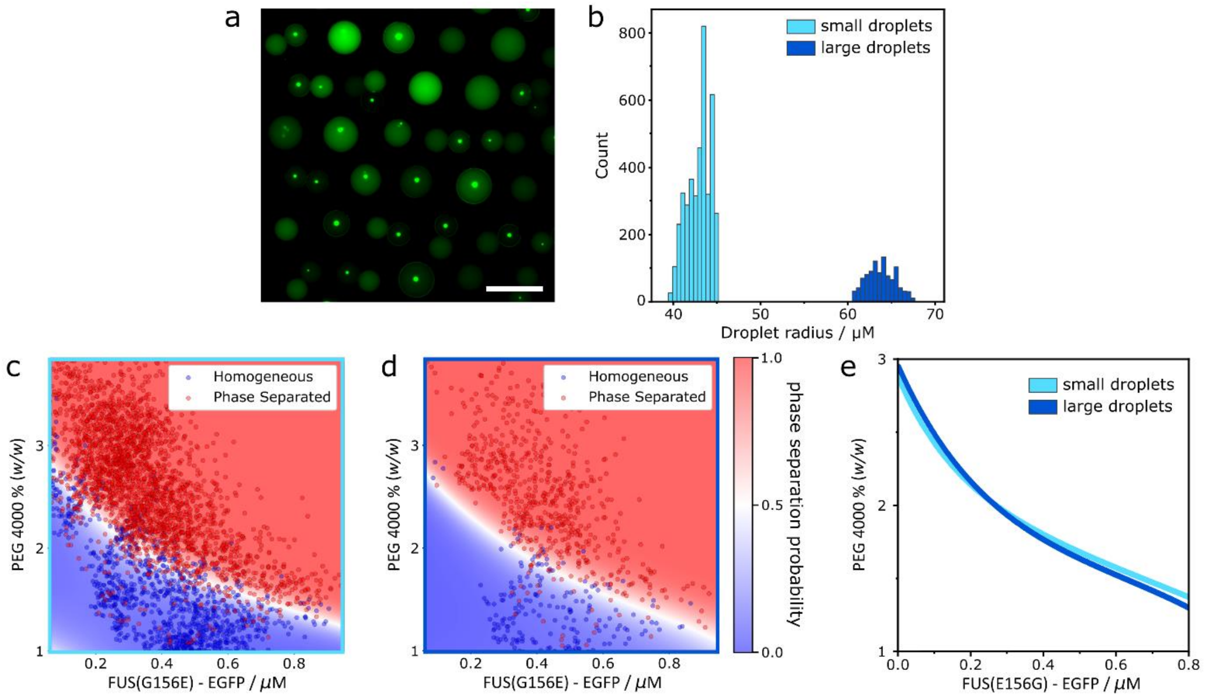


**Figure S5. Effect of droplet size on condensate phase behaviour. (a)** Epifluorescence microscopy images of trapped microdroplets with two different populations of droplet size. Shown is EGFP fluorescence corresponding to EGFP-tagged $FUS^{G156E}$. **(b)** Histogram of microdroplet size distribution showing a small ($r$ = 43 μm) and a large ($r$ = 65 μm) droplet population. **(c, d)** Phase diagram of EGFP-tagged $FUS^{G156E}$ vs. PEG concentration as obtained from experiments performed in small (panel c) and large (panel d) microdroplets. **(e)** Overlaying the computed phase boundaries obtained from small and large droplets showed negligible difference in phase behaviour between the two populations.

To investigate whether droplet size influences the observed phase behaviour, an experiment was performed in which a phase diagram for EGFP-$FUS^{G156E}$ and PEG was recorded using two different populations of droplet size (Figure S4(a)). Droplets with mean radius of 43 μm (small droplets) and 65 μm (large droplets) were produced by operating the droplet generator with oil flow rates of 300 μL/h and 60 μL/h, respectively (Figure S4(b)). Notably, this range of droplet size (3.5-fold difference in volume and 2.3-fold difference in surface area) is much larger than that produced during a conventional PhaseScan experiment, where droplet radii typically vary by <10%. Droplet imaging and analysis was executed as described in the Methods, with the data segregated between the two droplet sizes to afford phase diagrams for each of these droplet populations (Figure S4(c, d)). By overlaying the computed phase boundaries for the small and large droplets, it is apparent that no significant difference in phase behaviour between the two populations is observable (Figure S4(e)). Phase separation in the binodal region of phase-space occurs through a nucleation/growth mechanism, and phase separation could therefore be assumed to display volume and/or surface dependence as observed previously.[1] However, in the experimental procedure outlined here, droplets are assayed several minutes after generation and mixing. Therefore, we propose that each droplet microenvironment has sufficient time to reach chemical equilibrium, with the PhaseScan measurement invariant to droplet size over the range investigated here.

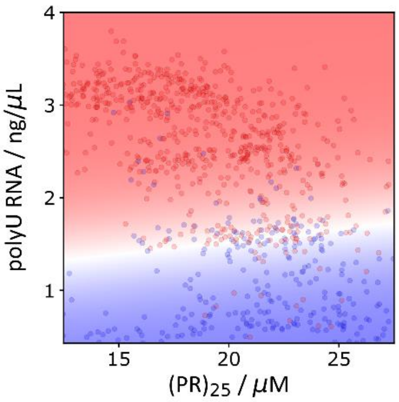


**Figure S6. PhaseScan characterisation of phase separation of (PR)$_{25}$ and polyU RNA.** To demonstrate the applicability of the PhaseScan assay to simple peptide systems as well as full-length proteins, we examined the phase behaviour of a dipeptide repeat system derived from the hexanucleotide repeat expansion in the chromosome 9 open reading frame 72 (C9orf72) gene, implicated in ALS.[2] The peptide consisted of 25 repeats of proline–arginine dipeptide (PR)$_{25}$. This type of peptide is well known to phase separate when mixed with negatively charged polymers, including single-stranded RNA. We assayed the formation of (PR)$_{25}$ with poly uridine (PolyU$_{100}$) RNA and observed RNA-dependent phase separation above 1.5 ng/μL RNA for ~20 μM (PR)$_{25}$.

#### Additional microdroplet generator designs

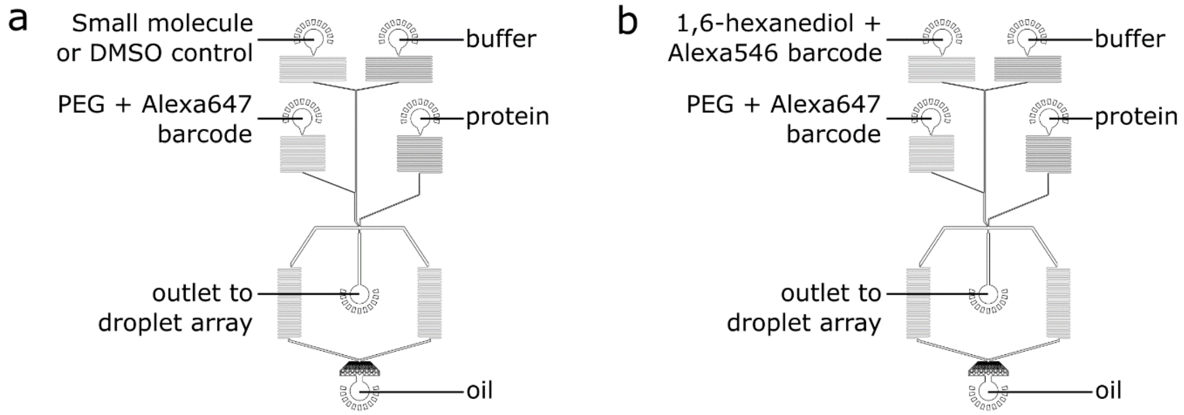


**Figure S7. Schematics of microfluidic devices for generation of microdroplets containing four aqueous components. (a)** Device design and input configuration for analysis of the effect of small molecules on phase separation (Figure 4, main text). **(b)** Device design and input configuration for generation of three-dimensional phase diagrams (Figure 4, main text).

#### Combined PhaseScan and droplet-shrinking methodologies

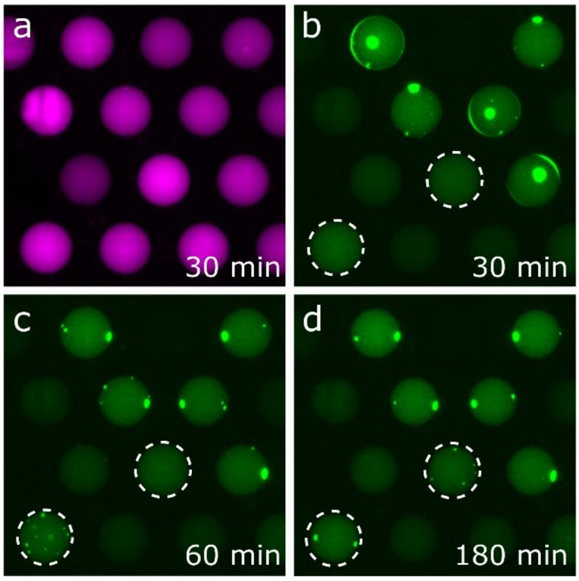


**Figure S8. Shrinkage of droplets following PhaseScan experiment. (a)** Epifluorescence microscopy image of Alexa647 dye encoding polyU RNA concentration 30 min after droplet trapping. **(b–d)** Images of FUS$^{G156E}$-EGFP fluorescence of phase-separated and homogeneous droplets 30, 60 and 180 min after droplet trapping, respectively. Droplets observed to transition from homogeneous to phase-separated regime are highlighted by dashed outline.

#### Supporting Materials and Methods

##### Protein production

FUS$^{G156E}$-EGFP expression and purification was adapted from Patel *et al.*[3] In short, recombinant protein production was performed in Sf9 insect cells (Expression Systems, Cat#94-001F) using the baculovirus system.[4] The protein was produced as a C-terminal EGFP fusion with an N-terminal maltose-binding protein (MBP) tag and a C-terminal hexahistidine ($His_6$) tag. Cells expressing MBP-FUS-EGFP$^{G156E}$-$His_6$ were harvested 72 h post-infection, centrifuged, and then resuspended in 50 mM Tris-HCl (pH 7.4), 1 M KCl, 5% (*w*/*v*) glycerol, 1 mM DTT, 10 mM imidazole supplemented with EDTA-free protease inhibitor cocktail set III (Calbiochem) and 0.25 U/mL benzonase (provided by the protein expression facility of the Max Planck Institute of Molecular Cell Biology and Genetics (MPI-CBG), Dresden). After cell lysis, using a shear homogenizer (Microfluidics), the protein was purified by immobilized-metal ion affinity chromatography (IMAC) using nickel-nitrolotriacetic acid (Ni-NTA) columns (Macherey-Nagel). The column was washed with lysis buffer. Elution was done with 50 mM Tris-HCl (pH 7.4), 1 M KCl, 5% (*w*/*v*) glycerol, and 500 mM imidazole. $His_6$ and MBP tags were proteolytically removed with 3C-$His_6$ preScission protease (provided by the protein expression facility of the MPI-CBG, Dresden). Protein was further purified by size-exclusion chromatography (SEC) using a Superdex 200 pg 26/600 column (GE Healthcare). Aliquots containing the protein were flash-frozen and stored at –80°C. The protein was stored in 50 mM Tris-HCl (pH 7.4), 500 mM KCl, 1 mM dithiothreitol (DTT), 5% (*w*/*v*) glycerol.

G3BP1-GFP expression and purification was carried out as described in Guillén-Boixet *et al.*[5] Briefly, recombinant $His_6$-GFP-G3BP1-MBP was expressed in and purified from insect cells (Expression Systems, Cat#94-001F) using a baculovirus expression system.[4] Following lysis (EmulsiFlex-C5, Avestin) in buffer containing 50 mM Tris-HCl (pH 7.5), 1 M KCl, 2 mM EDTA, 2 mM DTT and 1x EDTA-containing protease inhibitor cocktail (Roche), the protein was purified by affinity chromatography using amylose resin (New England Biolabs) to capture the protein via its MBP tag from the supernatant of the cell lysate. The sample was then subjected to IMAC using Ni-NTA resin (Qiagen). The column was washed with an EDTA-free lysis buffer containing 20 mM imidazole. The protein was subsequently eluted from the Ni-NTA column with 250 mM imidazole. $His_6$ and MBP tags were cleaved off with PreScission protease during an overnight dialysis step at 4$^{o}$C. The protein was further purified by SEC using a HiLoad 16/600 Superdex 200 pg (GE Healthcare) on an Akta Ettan system in 50 mM Tris-HCl (pH 7.5), 300 mM KCl, 1 mM DTT buffer. Aliquots containing the protein were flash-frozen and stored at –80°C.

NSP2 from Rotavirus A (strain RF) was produced as a C-terminal $His_6$-tagged protein (NSP2-$His_6$) in BL21(DE3) *Escherichia coli* transformed with a pET-28b-NSP2 construct, as previously described[6,7]. Expression was carried out at 24°C in Luria Bertani (LB) media supplemented with 1% (*v*/*v*) glucose, and cultures were induced with 0.5 mM IPTG once they reached optical density ($OD_{600}$) of 0.6, and harvested 14 h post-induction. Cell pellets were resuspended in 50 mM Tris-HCl (pH 8), 300 mM NaCl, supplemented with 100 µg/mL chicken egg lysozyme (Sigma), 0,5% Tween 20 and a complete protease inhibitor (Roche), followed by DNAseI treatment (10 µg/mL, Roche) for 15 min prior to clarifying the lysate by centrifugation for 20 min at 10,000 rpm at 4°C. Clarified lysate was loaded on a 5 mL His-Trap HP column (GE Healthcare), followed by a wash step and elution with 0.5 M imidazole, 20 mM HEPES-Na (pH 7.5). Eluted peak fractions were pooled, diluted with 10 mM HEPES-Na (pH 7.5) and purified by ion-exchange chromatography (IEX) step using a CaptoImpRes SP column (GE Healthcare). Eluted peak fractions were further resolved by SEC on a Superdex 200 10x300 GL column (GE Healthcare) pre-equilibrated with 25 mM HEPES-Na, pH 7.5, 150 mM NaCl. Purified protein aliquots were snap-frozen and stored at –80 °C for subsequent use. Labelling of the purified His-tagged NSP2 protein was achieved by pre-incubating 10 µM NSP2 with 1 µM Atto488-NTA dye (Sigma) for 5 min. Following incubation, the protein was immediately used.

Recombinant NSP5 from rotavirus A (strain RF) was produced as a N-terminal Strep-tagged protein in BL21(DE3) *E.coli* transformed with a pET-28b-NSP5 construct, as described.[8] Cultures (LB medium as above with NSP2) were induced with 1 mM IPTG, and expression was carried out for 6 h at 37°C. After enzymatic lysis as described above for NSP2, the protein was purified under denaturing conditions, followed by its refolding, as described.[9] Briefly, washed inclusion bodies were solubilized in 6 M guanidinium hydrochloride and the protein-containing fraction was then subjected to a refolding protocol following step-wise dialysis. After refolding, NSP5-containing fractions were further purified by IEX using a CaptoQ ImpRes column (GE Healthcare). Concentrated peak fractions were further resolved using SEC on a Superdex 200 10x300 column (GE Healthcare). Purified protein fractions were pooled, aliquoted and snap-frozen using liquid nitrogen, and stored at –80°C for subsequent use.

For SARS-CoV-2 Nucleocapsid (N) protein purification, baculoviruses were produced from a pOCC102-$His_6$-MBP-3C-Nucleocapsid construct, leveraging the FlexiBAC approach as previously described. The N coding sequence was derived from SARS-CoV-2 lineage B (GenBank: MN908947.3)[10] and subsequently codon-optimized for insect cell expression. Baculoviruses were then used in conjunction with an in-house Sf9 insect cell expression system (provided by the protein expression facility, MPI-CBG)[4]. Briefly, 0.5 L Sf9 cells (~$10^6$ cells $mL^{-1}$) were infected with 2% (*v*/*v*) baculoviral supernatant and subsequently grown for 72 h at 27°C and 85 rpm. Cell pellets were obtained by centrifugation at 1000 rcf for 5 min and routinely flash frozen in liquid nitrogen, and then stored at −80 °C. For cell lysis, Sf9 pellets were thawed and resuspended in 50 mL cold lysis buffer (1 M NaCl, 50 mM $Na_xH_xPO_4$, 20 mM imidazole, 5% (*v*/*v*) glycerol, 4 mM $MgCl_2$, 1 mM DTT, 1x complete protease inhibitor, 1 U $mL^{-1}$ DNase I, pH 7.4) and passed through an LM20 microfluidizer (15,000 psi, 4°C). Following ultracentrifugation (30,000 rpm, 4°C, 30 min), the obtained supernatant was passed through a 0.45 µm filter and subjected to a three-step FPLC purification on an ÄKTA pure 25 M chromatography system (Cytiva) at room temperature. Filtered cell lysates were first passed, at 5 mL $min^{-1}$, through the $Ni^{2+}$ NTA resin of a preequilibrated 5 mL HisTrap HP column (Cytiva). This was followed by a washing step with 40 mL imidazole wash buffer (150 mM NaCl, 50 mM $Na_xH_xPO_4$, 20 mM imidazole, 5% (*v*/*v*) glycerol, pH 7.4), and an imidazole gradient elution over 50 mL, reaching 300 mM imidazole, under otherwise identical buffer conditions. Fractions, of 1.5 mL each, were collected throughout and those with high absorbance intensities at 280 nm were analysed by SDS polyacrylamide electrophoresis and subsequently pooled. Following the concentration of pooled fractions, supplemented with additional 400 mM NaCl, 150 mM Arg-HCl (pH 7.4) and 300 mM trehalose to prevent aggregation, by repeated centrifugation (4,000 rcf, 25°C, 3 min) in a 30 kDa cut-off filter column to ~5 mL, purified proteins were digested using 500 µg of 3C protease (in-house purification; protein purification facility, MPI-CBG) in the presence of 0.5 mM DTT over 1 h at 25°C to remove the MBP solubility tag. Upon dilution of the reaction mixture to a final NaCl concentration of ~150 mM, the proteins were passed through the heparin-conjugated resin of a preequilibrated 5 mL HiTrap Heparin HP column (Cytiva). This was followed by a washing step with 40 mL heparin wash buffer (150 mM NaCl, 50 mM $Na_xH_xPO_4$, 5% (*v*/*v*) glycerol, pH 7.4) and a high salt gradient elution over 50 mL, ultimately reaching 1 M NaCl and pH 7.4, under otherwise similar buffer conditions. Fractions with putatively high protein content were again analysed and subsequently pooled. The pooled fractions were supplemented with additional 700 mM NaCl, 150 mM Arg-HCl (pH 7.4) and 300 mM trehalose to prevent aggregation, and concentrated by repeated centrifugation (4,000 rcf, 25°C, 3 min) in a 30 kDa cut-off filter column to ~2 mL. The concentrate was then passed through a 0.2 µm spin filter and resolved by SEC at a reduced flow rate of 0.5 mL $min^{-1}$ on a preequilibrated Superdex 200 Increase 10/300 GL column (~24 mL column volume; Cytiva), in SEC buffer (50 mM $Na_xH_xPO_4$, 300 mM NaCl, 5% (*v*/*v*) glycerol, 1 mM DTT, pH 7.4), which also constituted the final storage buffer. Following SEC, analysed and pooled fractions were again concentrated, as described above, in this case to ~200 µL. Employing a ND-1000 spectrophotometer (Thermo Scientific) the protein concentration was then determined at 280 nm (with a molar extinction coefficient ε ~43,900 $M^{-1}$ $cm^{-1}$ of the ~46 kDa CoV-2 N protein) and the extent of nucleic acid contamination evaluated based on the 260 to 280 nm absorption ratio (obtaining a value of ~0.56; a ratio of ≥0.7 indicating nucleic acid contamination). Subsequently, 5 µL aliquots were prepared, flash frozen in liquid nitrogen, and stored at −80 °C.

The $(PR)_{25}$ peptide, containing 25 proline–arginine repeats, was obtained from GenScript. N-terminally labelled PR25 was obtained by reacting the peptide with amine-reactive AlexaFluor546 (Sigma Aldrich). $(PR)_{25}$ experiments were conducted using a mix of 10% labelled and 90% unlabelled peptide.

**Device design and fabrication**

A standard lithographic process was applied to fabricate microfluidic devices.[11] The design of the device was drawn with AutoCAD (AutoDesk) and then printed on a photomask (Micro Lithography). The pattern of the mask was transferred by UV exposure[12] to a polished silicon wafer coated with a 50 µm thick layer of SU8-3050 photoresist (Microchem). Excessive SU-8 photoresist was removed using propylene glycol methyl ether acetate (PGMEA; Sigma). The wafer with SU-8 patterns (*i.e.*, master) was dried by blowing with nitrogen and baking at 95°C. The master was placed in a plastic petri dish and served as a mould for poly(dimethylsiloxane) (PDMS; Sylgard184, Dow Corning) casting. The PDMS base and crosslinking agent was mixed in a 10:1 ratio and polymerised by baking at 60°C for 2 h. The PDMS device was cut from the petri dish with a scalpel and cleaned by sonication in an ethanol bath for 15 min. Both the PDMS replica and the glass slide were activated in an oxygen plasma oven (30 s, 40% power, Femto, Diener Electronics) before bonding. The channels were treated with 1% (*v*/*v*) trichloro(1H,1H,2H,2H-perfluorooctyl)silane (Sigma) in HFE-7500 (Fluorochem) for 1 min, before being dried with nitrogen and heated on a hotplate at 95°C for 10 min.

**Operation of microfluidic devices**

Syringe pumps (neMESYS modules, Cetoni) were used to control flows of protein, buffer and phase separation trigger solutions (e.g., PEG) to the microfluidic device. The syringe (Hamilton 1710) and tubing (PTFE, 0.012"ID x 0.030"OD, Cole-Parmer) for the protein sample were prefilled with FC-40 oil, before the small working volume of protein sample (10–20 μL) was aspirated into the tubing. The aqueous flowrates were configured to vary automatically according to a pre-programmed flow profile with constant total flow rate of 60 μL/h. In this manner, variation of the concentration of the droplet components was achieved. FC-40 oil (containing 1.5% (*v*/*v*) fluorosurfactant) was introduced to the device at a constant flow rate of 50–120 μL/h depending on the phase separation system, with the use of high-viscosity PEG solutions requiring higher oil flow rate, for example. FC-40 oil (containing 1.5% (*v*/*v*) fluorosurfactant) was introduced to the droplet generator at a constant flow rate of 50 μL/h. The generated droplets were transferred via tubing to a separate microfluidic device (i.e., droplet catcher) which consisted of droplet traps arrayed in the roof of a flow chamber as described in Figure 1(a). After sufficient droplet generation time (5–6 min), the flow from all syringes was turned off and the oil syringe was connected to the droplet catcher via an intermediary microfluidic device, which is a simple channel that links the oil tubing to the droplet-catcher inlet tubing. The droplets were trapped in the wells of the droplet catcher at a constant oil flow rate of 300 μL/h, before excess droplets were flushed out of the device at an oil flow rate of 1000 μL/h. In some experiments (for example in acquisition of 3D phase diagrams), several droplet-catcher devices were linked in sequence to trap the desired number of droplets.

#### Image analysis

##### Calibration of droplet fluorescence intensity

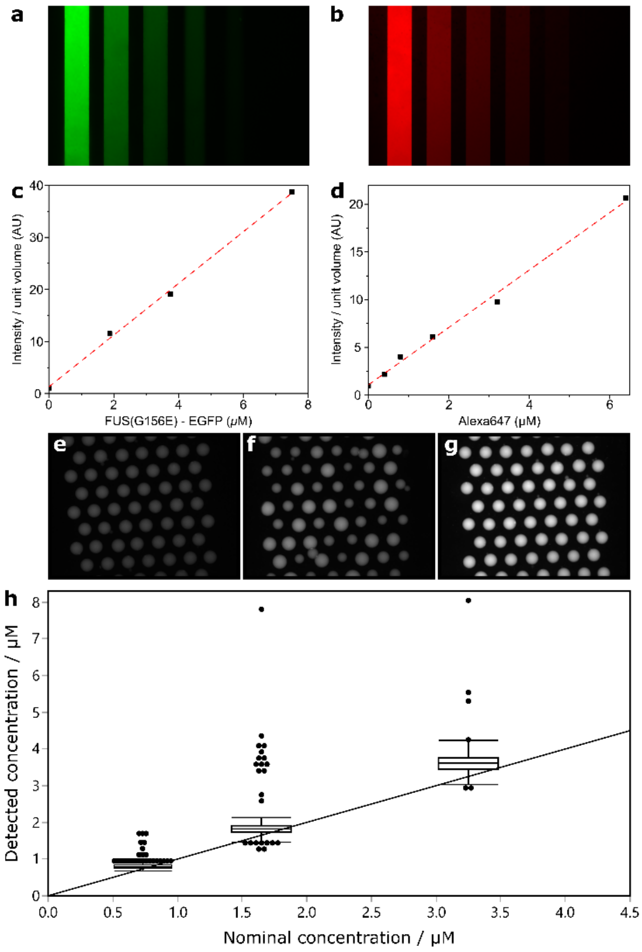


**Figure S9. Calibration of droplet fluorescence intensity. (a, b)** Representative epifluorescence microscopy images of known concentrations of EGFP-tagged FUS$^{G156}$ and Alexa647, respectively, contained in microchannels of known volume. **(c, d)** Calibration of EGFP-tagged FUS$^{G156}$ and Alexa647 concentration and calculated fluorescence intensity per unit volume, respectively. **(e, f, g)** Microdroplets containing Alexa647 at nominal concentrations of 0.73 µM, 1.65 µM and 3.25 µM, respectively. **(h)** Box plot of nominal Alexa647 droplet concentration and concentration as determined by the analysis program and intensity–concentration calibration procedure.

A calibration procedure was employed to convert the intensity per unit volume to the concentration of the corresponding barcode fluorophore (Figure S9) in order to allow each droplet to be accurately located in chemical space in the resultant phase diagram. The same fluorophore-containing solutions used in each experiment were injected into microchannels of known dimensions (cross sectional area typically 150 × 30 µm) and were imaged under the same conditions as the droplets (Figure S9(a, b)). From these images, the relationship between intensity and fluorophore concentration was determined on an experiment-by-experiment basis (to account for variation in pipetting, imaging, light source intensity etc.), as well as demonstrating the linear relationship between fluorophore concentration and fluorescence intensity (Figure S9(c, d)).

To demonstrate the efficacy of this approach, a control experiment was conducted where droplets containing known concentrations of Alexa647 fluorophore (as determined by UV-vis spectroscopy/NanoDrop) were produced, trapped, imaged and subjected to the analysis and calibration procedure (Figure S9(e, f, g)). The nominal fluorophore concentration present in each droplet population was observed to be in very good agreement with that determined by the analysis procedure (Figure S9(h)).

#### Detection and classification of droplets and condensates

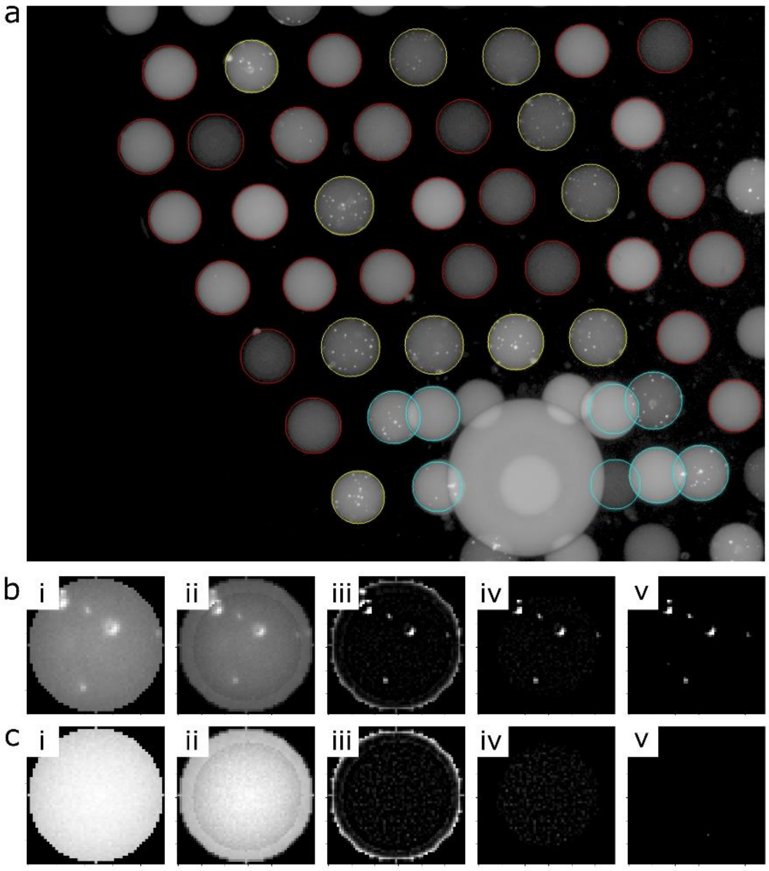


**Figure S10. Detection and classification of droplets and condensates. (a)** Yellow: phase separated droplets, red: homogeneous droplets, cyan: erroneously arrayed droplets. **(b, c)** Example droplet classification procedure for phase separated and homogeneous droplets, respectively. From left to right: (i) detected droplet, (ii) padding, (iii) convolution, (iv) cropping of the padded part, and (v) thresholding and identification of pixel clusters. For any droplet containing a cluster with >1 pixels, the droplet is considered as phase separated, otherwise mixed.

The collected images were analysed using a custom-written Python script (Figure S10). The raw images were processed to remove camera dark noise (Background) and flattened to correct non-uniform epifluorescence illumination by division with a calibration image taken of a homogeneous fluorescent solution (Illumination) according to $Processed\ Image = \frac{Image - Background}{Illumination - Background}$. The raw image was enhanced by taking the logarithm of the image, applying rolling ball background subtraction, taking the logarithm again, and thresholding the darkest 10% and brightest 10% pixels. Droplets were fitted as circles by finding the bright peaks in the image (Figure S8(a)). Any incomplete circles that are partly out of the image boundary as well as those smaller or larger than the radius thresholds were removed. Non-circular droplets or erroneous detections were removed by comparing a corresponding perfect sphere of same size and the image of the droplet, where the brightness multiplied by a proportional constant is used as the z-axis. The total intensity was calculated and normalised to afford intensity per unit volume (calculated using fitted diameter).

To distinguish droplets with condensates formed via phase separation and those which are well-mixed (Figure S10(b, c)), the droplet images were convoluted with an edge detection kernel after padding. The result image was made binary with a given threshold ratio of quartiles and medians. If there were at least two connected bright pixels, the droplet was classified as phase separated. Otherwise, the bright pixels were determined as noise and the droplet was labelled as well-mixed. The phase boundary was estimated using a support vector machine (SVM) algorithm with a radial basis function (RBF) kernel. The parameters of SVM were selected according to grid search scores. The parameter space was sampled as a 2D/3D mesh grid and predicted by the SVM model, which was then used to generate iso-boundary or iso-surface as the phase diagram boundary.
